# Large scale similarity search across digital reconstructions of neural morphology

**DOI:** 10.1101/2021.12.17.473026

**Authors:** Bengt Ljungquist, Masood A. Akram, Giorgio A. Ascoli

**Affiliations:** Center for Neural Informatics, Structures, & Plasticity and Bioengineering Department, George Mason University, Mail Stop 2A1, 4400 University Dr, Fairfax, VA, United States of America

**Author notes:** Corresponding author. *E-mail address:* (Giorgio A. Ascoli).

**Keywords:** Neuronal Morphology, Principal Component Analysis, Neuroinformatics, Similarity search, Software as a Service

## Abstract

Most functions of the nervous system depend on neuronal and glial morphology. Continuous advances in microscopic imaging and tracing software have provided an increasingly abundant availability of 3D reconstructions of arborizing dendrites, axons, and processes, allowing their detailed study. However, efficient, large-scale methods to rank neural morphologies by similarity to an archetype are still lacking. Using the NeuroMorpho.Org database, we present a similarity search software enabling fast morphological comparison of hundreds of thousands of neural reconstructions from any species, brain regions, cell types, and preparation protocols. We compared the performance of different morphological measurements: 1) summary morphometrics calculated by L-Measure, 2) persistence vectors, a vectorized descriptor of branching structure, 3) the combination of the two. In all cases, we also investigated the impact of applying dimensionality reduction using principal component analysis (PCA). We assessed qualitative performance by gauging the ability to rank neurons in order of visual similarity. Moreover, we quantified information content by examining explained variance and benchmarked the ability to identify occasional duplicate reconstructions of the same specimen. The results indicate that combining summary morphometrics and persistence vectors with applied PCA provides an information rich characterization that enables efficient and precise comparison of neural morphology. The execution time scaled linearly with data set size, allowing seamless live searching through the entire NeuroMorpho.Org content in fractions of a second. We have deployed the similarity search function as an open-source online software tool both through a user-friendly graphical interface and as an API for programmatic access.

## 1. Introduction

Since the dawn of neuroscience, with the elegant drawings of Ramón y Cajal [1], it has been known that the branching morphology of neuronal and glial (henceforth referred to as neural) arbors underlie their functional role in the nervous system. Specific neuron types often have characteristic dendritic or axonal trees, as exemplified by the iconic structures of cerebellar Purkinje cells and cortical chandelier cells, respectively [2]. Multifarious analysis approaches have been developed both to establish common morphological characteristics within neural types and to compare structural differences between types [3]. Quantitative representations of tree morphologies harnessed, for example, different morphometrics statistics [4]–[7] and spatial density maps [8], [9]. Arbor topology was captured through sequence representation [10] or using a local alignment between pairs of neurons via dynamic programming [11]. More recently, algebraic methods based on persistent homology allowed to quantify branching distribution into vector metrics [12], [13].

It should be possible to leverage these characterizations to find similar morphologies from a large set, given a neuronal or glial reconstruction of interest as the query vector. To this purpose, the NBLAST algorithm combined cell position with local geometry [14]. A different method for global search employed an asymmetric binary coding strategy based on the maximum inner product [15], while encoding of morphology with hashing forests has shown promising performance on large datasets [16]. Another variant is to query sub-structures of the neuron with graph representations of the (sub-)trees [17] or using structure tensors and expanding this field via gradient vector flow [18]. A conceptually related problem is to compare two morphological reconstructions of one and the same neuron, such as when benchmarking an automated tracing software against the gold standard of expert manual proofreading [19].

For the past 15 years, NeuroMorpho.Org enabled sharing of 3D reconstructions of neural morphology [20], with the latest major release, 8.1, accruing 150,000 digital tracings from 78 species, 1330 cell types, 381 brain regions, and 774 contributing labs. The growth rate of this repository has continuously increased due to a combination of more efficient reconstruction techniques, greater willingness of the neuroscience community to share, and rising open data expectations from funding organizations and scientific publishers [21]. In parallel, the NeuroMorpho.Org internal processing pipeline evolved into a micro-service based architecture, reducing the time from deposition to publication from months to weeks [22]. Within the data standardization workflow, it must be verified that each submitted reconstruction is not a duplicate of any previously submitted specimen. This step, equivalent to an extreme case of similarity matching, used until recently Pearson correlation of morphometric data. However, such method was slow for larger data sets, prompting the development of the more efficient alternative described in this work.

To our knowledge, no readily available *Software as a Service* method provides precise and fast search of neural reconstructions solely by morphological similarity for larger data sets (>100k reconstructions). Therefore, we engineered a combination of the state-of-the-art software Facebook AI Similarity Search, or FAISS [23], with L-Measure statistical morphometrics [6] and persistence vectors descriptors [12]. We report that this system enables efficient and massive parallel searches, works flexibly with both global (whole cell) and local (parts of the cell) neural arbors, and is highly effective at duplicate detection. Moreover, we investigated the impact of dimensionality reduction using PCA (principal component analysis) in terms of precision and speed. Lastly, we present the new NeuroMorpho.Org functionality enabling users to carry out similarity searches through an intuitive graphic interface or an Application Programming Interface (API). All code is released open source to ensure maximum reproducibility and encourage further community development [24].

## 2. Materials and Methods

NeuroMorpho.Org provides for each reconstruction 21 morphometrics, for example, total length, average bifurcation angle, and the number of branches (for a complete list, see http://NeuroMorpho.Org/myfaq.jsp?id=qr4#QS3), as derived by L-Measure [6]. These metrics are calculated both for the entire cell, referred to as summary morphometrics, as well as separately for each structural domain, for example only the axon, and are then referred to as detailed morphometrics. For summary morphometrics, the dimensionality is thus 21, while for detailed morphometrics it is a multiple of 21 depending on the number of distinct structural domains, up to a maximum of 63 for a neuron with apical and basal dendrites as well as an axon.

In addition to the summary and detailed morphometrics, NeuroMorpho.Org also stores 100-dimensional persistence vectors quantifying the branch distribution by algebraic homology [12]. We have modified the original C++ and Java open-source code [12] to run on Linux and added a Python wrapper to make the software callable as a service. The source code of this modified software is available open source at https://github.com/NeuroMorpho/swc2pvec, and the service is briefly described and callable as an API at http://cng-nmomain.orc.gmu.edu/swp2pvec (for a complete API description, please see the GitHub page).

NeuroMorpho.Org also provides rich metadata for all reconstructions [25], including both non-numerical (e.g., species and brain region) and numerical specifications (e.g., slice thickness and objective magnification). For purposes of similarity search, we hashed and normalized non-numerical metadata values into a numerical representation, resulting in a 29-dimensional vector.

FAISS (Facebook AI Similarity Search) is a fast and memory-efficient similarity search software developed by the company Meta, formerly Facebook [23]. To generate an index of distances (similarity) from the input data, FAISS offers different methods trading off properties such as memory usage vs speed. We have consistently used a flat L2 index in our implementation. The index may then be queried using a vector of the same dimensions as the indexed data. We have also normalized all data before building the index. The resulting similarity is the L2 normalized distance between the two vectors, ranging from −1 to 1, with −1 meaning they are maximally dissimilar, and 1 that they are identical. In addition to FAISS, we have implemented for benchmarking purposes a traditional Pearson correlation for similarity search of a single cell at a time (non-parallel). We have also tested a combination of the two, FAISS similarity multiplied by Pearson correlation, which thus also can only handle searches with one neuron at a time. Pearson correlation was calculated in real-time, as pre-calculation would take for the current database 130 GB of memory, far exceeding the available RAM. The RAM footprint of a typical similarity search in our implementation stayed around 2-3 GB.

The similarity search software was written in Python, including NumPy and Flask in addition to the FAISS Python library, and deployed as a Docker service using Ubuntu Linux 18.04 LTS as the operating system on a virtual machine hosted at the data center of George Mason University’s Office of Research Computing. We created the following data vectors for each neuron to build a similarity search, with dimension m as indicated: 1. Summary morphometrics (m=21); 2. Detailed morphometrics (m=21, 42 or 63, that is, 21 per structural domain present); 3) Persistence vectors (m=100); 4) 1 combined with 3 (m=121); 5) 2 combined with 3, (m=121, 142 or 163); and 6) Binary metadata comparison (m=29).

For the above morphometric data sets, we applied PCA by calculating the eigenvectors (principal components) and correspondent eigenvalues of the covariance matrix and sorting them in order of falling ratio of explained variance. We adopted the broken stick method for determining how many principal components to include as base vectors [26]. This method advocates stopping when the ratio of explained variance of the next principal component falls below what could be expected if performing the PCA on white noise, that is 1/m, where m is the dimension of the original data. For the selected principal components, we calculated the sum of explained variance as a measure of their information content.

A similarity search user interface was implemented in JSP and integrated with NeuroMorpho.Org, where, starting from any cell page of interest, users may search for similar cells choosing any of the 6 methods described above. As with other NeuroMorpho.Org search functionalities, the SWC files of the found neurons or glia can be immediately visualized or saved for separate downloading. Users may also select to apply PCA and whether to utilize FAISS, Pearson correlation, or their scalar product. The graphical user interface in turn calls an underlying API, which is also independently machine-accessible (https://github.com/NeuroMorpho/similarity-search).

In order to assess the capability of the software to rank morphological similarity, we performed a visual evaluation. Similarity search was performed on a representative set of 100 reconstructions from NeuroMorpho.Org selected so as to cover the broad span of species, brain regions, cell types, and experimental methods in the database. The search used summary morphometrics plus persistence vector as data, dimensionality reduction using PCA, and combined FAISS and Pearson correlation as index. The top six most similar cells were then compared with six randomly selected cells relative to the original reconstruction, and their similarity scores calculated. Furthermore, we generated a histogram of the similarity scores from each target cell to all other cells in the database.

To evaluate performance scaling for parallel searches, that is searching simultaneously through the entire NeuroMorpho.Org content for entries similar to many cells, we used persistence vector plus summary morphometrics (m=121) and compared that with the same after dimensionality reduction by PCA (m=8). For the two approaches, we varied the number of parallel searches from 200 cells to 4800 in 200 increments and measured the similarity search execution time. This was performed using FAISS as Pearson correlation cannot handle parallel searches.

We then evaluated the ability to find data duplicates of the different morphometric indices, with and without PCA applied, thus comparing six different methods. Detailed measurements were not used as they cannot compare cells of different structural domains. Likewise, we did not use metadata similarity for the purpose of duplicate detection, as cells often share the same metadata when they are contributed from the same lab to the database. We compared the efficiency of these six methods in finding duplicates by calculating the false negative rate (FNR or miss rate) and false positive rate (FPR or fall-out) for duplicates detection. The trial for this detection utilized the 131,960 reconstructions from the latest major release of NeuroMorpho.Org that contained duplicates (v. 8.0), since those have been cleared out as of v. 8.1. Starting from a set (1000 cells) of potential duplicates identified using all three descriptors as well as an archive-by-archive inspection, we manually generated a list of 235 true duplicates by confirming through visual inspection and examination of the source files and related annotations provided by the original contributors. For all 131,960 reconstructions, we then applied parallel FAISS similarity search for an all-against-all duplicate detection using all six methods separately. This generated a list of the 10 most similar cells for each method and cell. If any of these had a similarity score >=0.9999, it was considered a potential duplicate. This was then compared against the list of true duplicates and, if confirmed, considered a true positive, otherwise a false positive. If a cell pair was on the list as a true duplicate, but fell below the similarity threshold, it was considered a false negative. Lastly, cell pairs that neither were on the true duplicate list nor passed the similarity threshold were considered true negatives. We also evaluated impact to FPR and FNR when setting the required similarity score to >=0.99999 or >=0.999.

The implementation is provided open source at the NeuroMorpho.Org GitHub account: https://github.com/NeuroMorpho/similarity-search. Database credentials have been removed, as direct database access is not provided to outside users for security reasons, but all data used for similarity search is provided as Python pickle files (*.pkl). It is also possible to download the application as a Docker container image, including the pickle files: https://hub.docker.com/repository/docker/neuromorpho/sis.

## 3. Results

The visual evaluation of representative cells demonstrated that the similarity search selects visually similar neurons when compared to random neurons (Figure 1). All other inspected cells showed analogous results in terms of the ability to find visually similar cells. Moreover, we noted that the distribution of similarity scores for an individual cell against the whole database was shaped differently (unimodal or bimodal, right-tailed or left-tailed etc.) for distinct cell types, such as projection neurons, interneurons, and glia.

**Figure 1.**
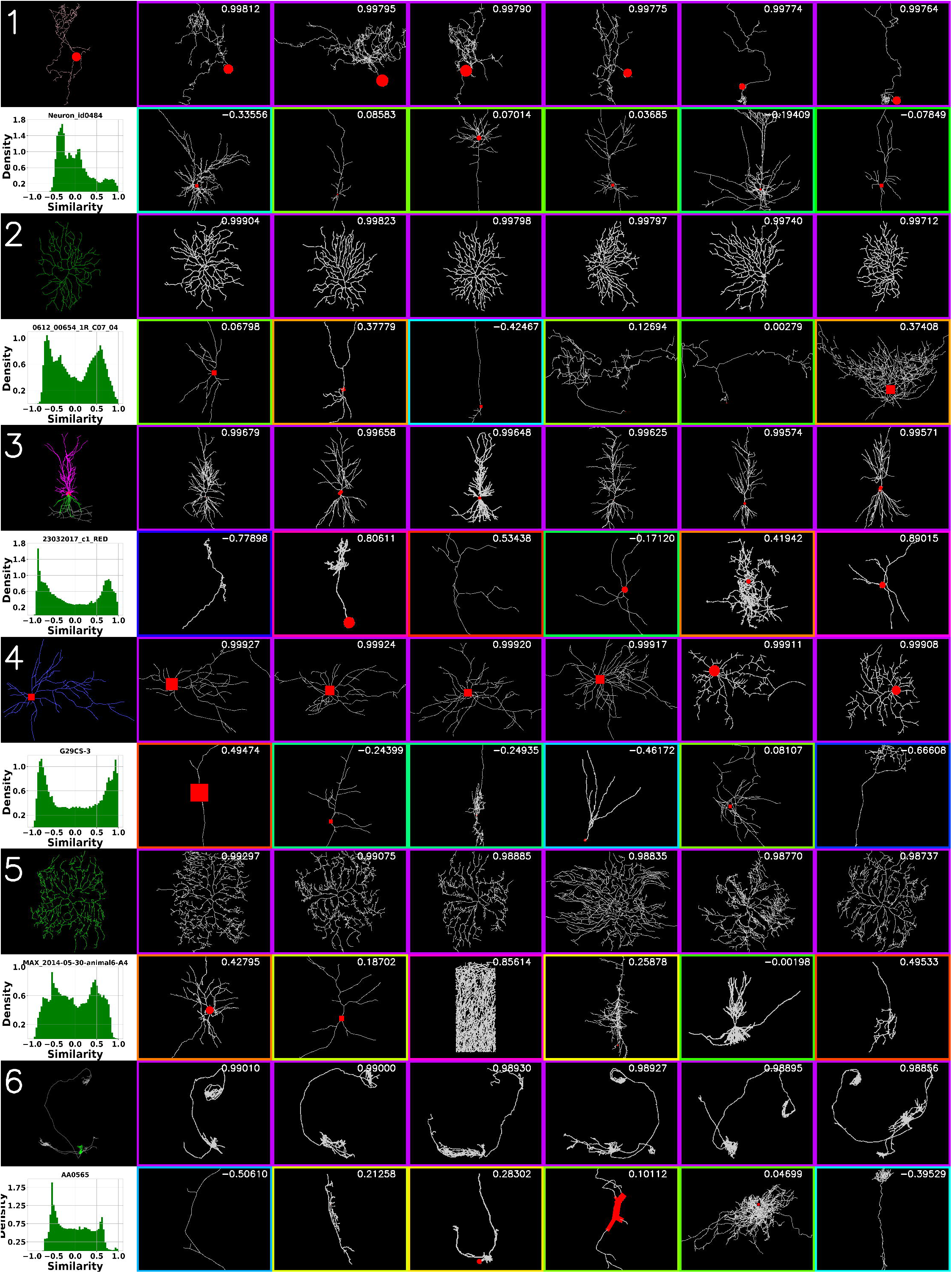
Similarity search results compared to random controls for six representative cells from NeuroMorpho.Org: 1) zebrafish main olfactory bulb interneuron [28] (NeuroMorpho identifier: NMO_93878); 2) mouse retinal ganglion cell [29] (NMO_128003); 3) mouse hippocampal pyramidal cell [30] (NMO_94734); 4) rabbit corpus callosum astrocyte [31] (NMO_53207); 5) drosophila melanogaster peripheral nervous system sensory neuron [32] (NMO_76528); and 6) mouse hippocampal projection cell [33] (NMO_121747). Top left figure in each row is the search target. The histogram directly below each target cell plots the similarity of that cell against all others in the database. Immediately to the right of each target cell are the six most similar cells found by the search in falling order of similarity score (indicated in top right corner of each cell). On the second row, to the right of the histogram, six random cells are shown with their similarity score in the top right corner as well. The color of the frame of each cell designates its similarity, with purple the most similar (score 1), green neutral (score 0), and blue the least similar (score −1). The original target cells are colored with respect to structural domain, while all other cells are greyed out to avoid visual bias.

The implemented microservice-based similarity search allows for fast and efficient search both for machine and human users. In particular, starting from any “cell page” (e.g., http://neuromorpho.org/byRandom.jsp), NeuroMorpho.Org users can search for similar morphologies after selecting preferred search parameters (Figure 2). Choices include the similarity implementation to use (FAISS and/or Pearson correlation), the numerical descriptors (summary morphometrics, persistence vectors, their combination, detailed morphometrics, or metadata), whether or not to apply PCA, and the result size (10, 25, 50, or 100 most similar hits). Results are delivered within a fraction of a second in all cases. The underlying API has the same parameter options for programmatic access.

**Figure 2.**
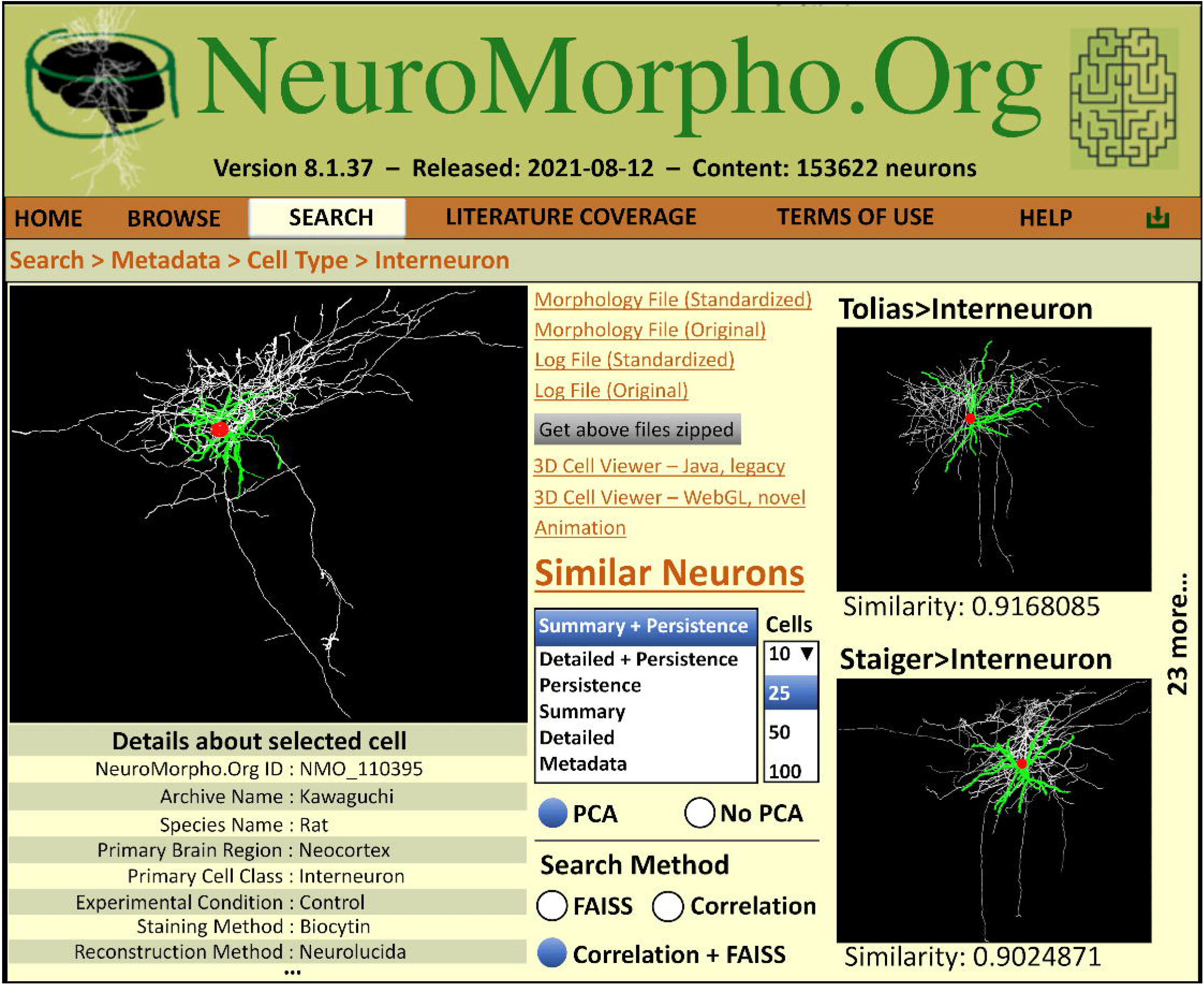
Similarity search with the graphical user interface. Starting from any cell page as in the illustrated example of an interneuron [34] (left), users can access the new Similarity Search functionality and select the desired options (center, bottom). The results are displayed in order of similarity scores among the entire content of the database (right). The illustration has been adapted slightly from the current user interface look for optimal display.

Search response time was linear with the number of parallel searches: 0.15s for 1000 cells, 0.3s for 2000, and 0.6s for 4000. The time difference between summary morphometrics plus persistence vectors with PCA (9 dimensions) and no PCA (121 dimensions) applied was constant with a mean of 0.05s.

The broken stick method for selecting principal components of the three descriptors resulted in different results for each (Figure 3). For the persistence vectors, summary morphometrics, and their combination, the number of eigenvalues greater than the cut-off limit were 7, 4, and 8, with a sum of explained variance of 96.5%, 91.3%, and 97.1% respectively.

**Figure 3.**
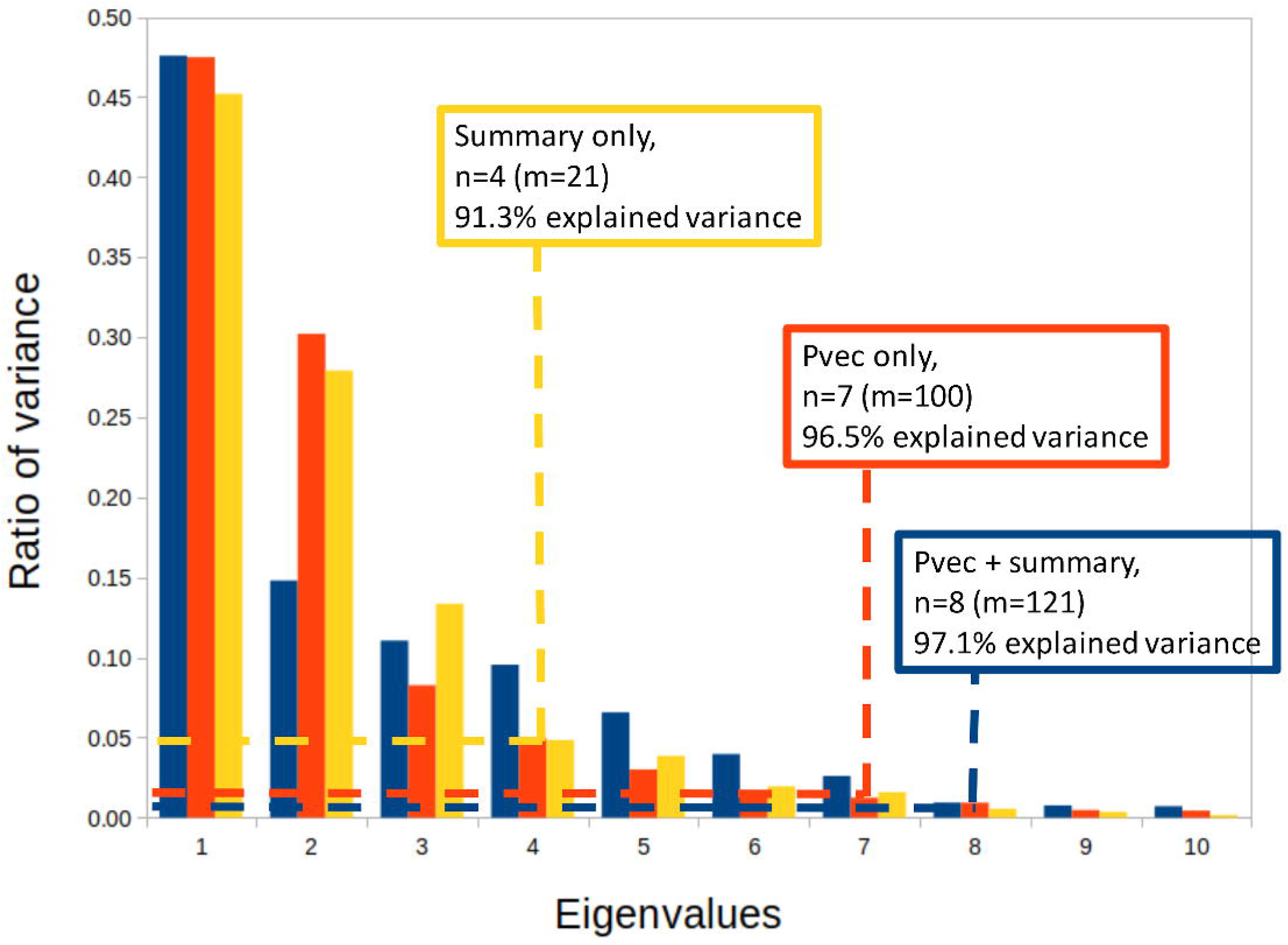
Ratio of variance of each eigenvalue for the three different descriptors used in similarity search. Dashed lines represent the calculated cut-off *n* using the broken stick method (>1/m where m is the original dimension).

Evaluation of the duplicate detection test showed that combining summary morphometrics and persistence vectors at a similarity threshold of 0.9999 is effective for finding the occasional repeated reconstructions (**Table 1**). Notably, persistence vectors alone performed worst, while summary morphometrics performed better but still far worse than the combined method both in terms of false positive rate (FPR) and false negative rates (FNR). Applying PCA improved both FPR and FNR for persistence vectors, while performing worse for FNR using summary morphometrics and slightly worse for the combined method, but substantially better for FPR for both. Using a higher similarity threshold of 0.99999 improved FPR slightly to 0.18% but worsened FNR dramatically to 29.87% for the combined method with PCA. Conversely, reducing the threshold to 0.999 completely eliminated false negative but yielded an unacceptably high FPR of 7.4%.

**Table 1.** Duplicate detection false positive rate (FPR) and false negative rate (FNR) for the different methods at a threshold similarity of 0.9999, expressed as percentages with standard deviation. A total of 235 true duplicates were presented in the set of 131,960 cells used for the statistics calculation.

| Method | Full - FPR | PCA - FPR | Full - FNR | PCA - FNR |
| --- | --- | --- | --- | --- |
| Persistence vectors | 35.37% ± 0.13% | 19.91% ± 0.11% | 24.68% ± 0.12% | 17.02% ± 0.10% |
| Summary Morphometrics | 17.34% ± 0.10% | 6.66% ± 0.07% | 2.55% ± 0.04% | 9.79% ± 0.08% |
| Persistence vectors + summary morphometrics | 1.34% ± 0.03% | 0.49% ± 0.02% | 0.43% ± 0.02% | 0.85% ± 0.03% |

## 4. Discussion

In this work, we introduced and evaluated a similarity search among neuronal and glial reconstructions based on three numerical descriptors of morphology. When combined with the Facebook AI Similarity Search (FAISS) software, the result is fast, efficient, and precise. In addition to sharing the code freely for further development and standalone implementations, we deployed this new function in the publicly available database NeuroMorpho.Org both as a userfriendly graphical interface and as API. Users can perform seamless similarity searches against hundreds of thousands of cells easily and quickly. Common applications include finding reconstructions similar to a target archetype of interest or identifying occasional duplicates across large data sets. The results obtained with FAISS are qualitatively similar to those yielded by traditional Pearson correlation, but FAISS additionally enables massive parallel searches. This allows the programmatic ranking of similarities for thousands of reconstructions simultaneously with results in fractions of a second. Such a performance, which to our knowledge has not been previously achieved, may facilitate ever more powerful unbiased classification of neural morphology [27].

A visual inspection of the similarity results showed that the method has a strong ability to find similar reconstructions to a representative variety of target morphologies when compared to a random selection. Interestingly, the distributions of the similarity scores against the entire NeuroMorpho.Org database varied substantially from one target cell to another. We observed that cells with similar metadata tended to display comparable similarity distributions, but a detailed investigation of these distributions remains a subject for future study.

We evaluated the information content of the three morphological descriptors: persistence vectors, L-Measure summary morphometrics, and the combination of the two, by studying the explained variance after PCA application. Persistence vectors combined with summary morphometrics constitute the most information rich descriptor, with a reduction from 121 to 8 dimensions retaining over 97% of the variance. The summary morphometrics, with PCA dimensionality reduction from 21 to 4, only retained 91.3% of the variance, while the persistence vectors, with dimensionality reduced from 100 to 7 post-PCA, retained 96.5% of the original variance. Our interpretation is that summary morphometrics alone is a noisier descriptor but delivers superior precision for similarity search when combined with persistence vectors.

Execution time performance scales linearly with a growing number of cells in a parallel query. However, larger dimensions of each query vector have a comparatively smaller impact, as the similarity with no PCA applied (m=121) was only marginally slower than the similarity search with PCA applied (m=8). It should be noted that we have used the CPU implementation of FAISS on standard server hardware, with no GPU (Graphics Processing Unit) support. This was sufficient for our purposes, as the similarity search in most cases returned results within a fraction of a second. The search is expected to execute substantially faster when applying GPU acceleration. The method is therefore likely suitable also for exceptionally large data sets including whole-brain data sets of millions of neural reconstructions.

Using persistence vectors and summary morphometrics combined as a descriptor constitutes in our experience the strongest similarity search function. This same combination also provides the most effective duplicate detection test and is superior to the persistence vectors and summary morphometrics used separately as query vectors. We also noticed that applying PCA generally yields improved results for all three methods investigated when considering both false positive and negative rates. Therefore, we recommend the combined method with PCA as the default setting for NeuroMorpho.Org similarity searches, although different options should be explored when working with defined subgroups of cells, such as only microglia, only cultured cells, only long axonal projections, or only invertebrate neurons.

Similarity search is available for both human users through the NeuroMorpho.Org graphical interface as well as for programmatic usage as an API, therefore allowing all users to discover morphologically similar neurons efficiently. The now fully automated microservice-based data ingestion pipeline of NeuroMorpho.Org [22] checks for duplicate reconstructions before ingestion as a part of the data quality assurance. This is necessary as reconstructions may unintentionally be resubmitted if they are used in different studies together with novel reconstructions. Lastly, this work highlights NeuroMorpho.Org as a mature database, which provides the necessary data diversity and quality required for a precise similarity search.

## Acknowledgements

The authors thank Praveen Menon, Navy Merianda, and Pruthvik Narayanaswamy for their contributions in development of a prototype for similarity search. This research was supported in part by NIH grants U01MH114829, R01NS39600, and R01NS08608

